# Identify Alzheimer’s disease subtypes and markers from multi-omic data of human brain and blood with a subspace merging algorithm

**DOI:** 10.1101/2025.04.30.651565

**Authors:** Ziyan Song, Xiaoqing Huang, Asha Jacob Jannu, Travis S. Johnson, Jie Zhang, Kun Huang

## Abstract

Identifying Alzheimer’s disease (AD) subtypes is essential for AD diagnosis and treatment. We integrated multiomics data from brain tissues of the ROSMAP and MSBB studies using a subspace merging algorithm and identified two AD patient clusters with notable cognitive and AD pathology differences. Analysis of differentially expressed genes (DEGs) in brain and blood samples pinpointed the LDLR gene as a potential blood biomarker linked to brain gene expression changes. Furthermore, we conducted PheWAS analysis on All of Us Project’s EHR and WGS dataset for 105 eQTLs associated with the DEGs and revealed significant associations between these eQTLs and several phenotypes, shedding light on potential regulatory roles of these genes in diverse physiological processes. Our study successfully integrated multiomics data and proposes LDLR as a candidate blood biomarker for AD subtyping. The identified phenotypic signatures provide valuable insights on molecular mechanisms underlying AD heterogeneity, paving the way for personalized AD treatment.

## Introduction

Alzheimer’s disease (AD) is characterized by a high degree of heterogeneity, encompassing diverse brain pathological changes, progression trajectories, and a multitude of risk factors among affected individuals[1]. Utilizing data-driven approaches for subtyping AD holds the promise for elucidating its etiology and profoundly improving AD diagnosis, treatment, and disease management strategies. This project endeavors to delineate subtypes of AD by leveraging integrative analyses of multi-omics data derived from postmortem brain and blood samples of AD patients.

Integration of multi-omics data for disease subtyping has been widely studied in diseases such as cancers. There are multiple challenges with multi-omics data integration including high-dimensionalities of the omics, different numerical ranges between data types, and strong relationships between data features (e.g., genes and proteins). These challenges make it challenging to integrate the features from all omics data for downstream subtyping analysis, which is often based on unsupervised learning methods. To address these challenges, several integrative approaches have been proposed. One strategy involves constructing patient similarity networks for each omics modality and merging them into a unified consensus network. Similarity Network Fusion (SNF) adopts this approach and has been widely applied to disease subtyping, including cancer and neurological disorders [2]. Another representative method is Multi-Omics Factor Analysis (MOFA), which models shared and modality-specific latent factors using a Bayesian framework. MOFA is particularly useful for exploring the underlying sources of variability across heterogeneous omics layers [3]. Similarly, iClusterPlus leverages a joint latent variable model and likelihood-based inference to simultaneously integrate multiple omics datasets with different distributions, such as Gaussian or binomial, and has been applied successfully to cancer subtype discovery [4]. In this study, we employ an advanced subspace merging algorithm to conduct unsupervised clustering analysis on AD patients, integrating gene expression, proteomics, and DNA methylation data from two prominent studies: Mount Sinai Brain Bank (MSBB)[5] and the Religious Orders Study (ROS) and Memory and Aging Project (MAP)[6]. Originally developed for cancer applications, the subspace merging algorithm demonstrated superior performance compared to SNF in benchmarking studies. Unlike MOFA, or iClusterPlus, our method takes a geometric perspective by aligning low-dimensional subspace representations of each omics data type on a Grassmann manifold. This design preserves modality-specific structure while capturing shared patient-level patterns, thus providing a flexible and mathematically grounded framework for discovering robust molecular subtypes.

Additionally, we aim to translating the subtypes discovered using brain tissues to blood samples, which can lead to discovery of blood markers in live patients. To achieve this, we incorporate RNA-Seq gene expression data collected from blood samples of matched patients within the ROSMAP study, expanding our analytical scope to peripheral biomarkers. Specifically, we performed differential expression analysis, enrichment analysis, and pathway analysis to establish potential connections between DEGs identified from brain samples and those highlighted from blood samples.

Furthermore, we investigated whether clinical phenotypes are associated with the expression levels of the discovered brain and blood markers. Such phenotypes may serve as early indicators or markers to predict AD risks. To test this notion, we extend our investigation by conducting a phenome-wide association study (PheWAS)[7] using comprehensive genomics data from the *All of Us* Research Project which contains both genotype data and extensive clinical records for a large heterogeneous cohort of adults covering different sexes and races. We then used established eQTLs[8] to identify genetic variants associated with the expression levels of the DEGs and then explored the association between the genetic variants of these eQTLs and a wide variety of phenotypes in the All of Us cohort.

The identification of blood-based biomarkers holds significant promise for advancing AD research and clinical practice, offering potential advancement for enhancing diagnostic accuracy, treatment efficacy, and preventive strategies. By exploring the molecular underpinnings of AD heterogeneity and establishing connections between brain and peripheral biomarkers, our study contributes to the growing body of knowledge aimed at improving the understanding and management of AD.

## Data Used in this Study

Three types of omics data, namely gene expression, DNA methylation data, and proteomics data, were obtained from the Alzheimer’s Disease Knowledge Portal for both the ROSMAP and the MSBB studies. In the case of ROSMAP, additional Monocyte RNA-seq data collected from matched AD patients were also integrated into the analysis. This comprehensive dataset included 79 AD patients with matched gene expression[9], DNA methylation[10], and proteomics[11] data from the dorsolateral prefrontal cortex (DLPFC) of autopsied brains. Additionally, a subset of 28 patients had additional gene expression data from Peripheral Blood Mononuclear Cells (PBMC) samples included in the analysis. Furthermore, for the MSBB study, multi-omics data from 83 AD patients were incorporated, including RNA-seq data collected from four distinct Brodmann areas: areas 22 and 36 from the temporal lobe, and areas 10 and 44 in the frontal lobe of the brain. Furthermore, Isobaric Multiplex Tandem Mass Tag (TMT) labeled proteomics and DNA Methylation array data from matched patients were included. Clinical data for all patients were also included in the follow-up downstream analysis. Detailed information regarding these datasets is provided in Table 1.

**Table 1.** Summary of Dataset Characteristics: Matched RNA-Sequencing data, DNA methylation data, and proteomics data were obtained from the dorsolateral prefrontal cortex (DLPFC) of 79 AD patients from the ROSMAP study, with an additional subset of 28 patients providing mRNA expression data from peripheral blood mononuclear cells (PBMC). Additionally, matched RNA-seq, DNA methylation, and proteomics data were collected from four distinct brain regions (Brodmann areas 10, 22, 36, and 44) in the MSBB dataset.

| Study | Omics Data Type | Number of Samples | Number of features |
| --- | --- | --- | --- |
| ROSMAP | mRNA DLPFC | 79 | 15582 |
|  | mRNA PBMC | 28 | 18625 |
|  | DNA Meth | 79 | 54596 |
|  | Proteomics | 79 | 5211 |
| MSBB | mRNA BM10 | 83 | 17401 |
|  | mRNA BM22 | 83 | 17401 |
|  | mRNA BM36 | 83 | 17401 |
|  | mRNA BM44 | 83 | 17401 |
|  | DNA Meth | 83 | 2608 |
|  | Proteomics | 83 | 12147 |

We conducted all integrative analyses separately within each cohort, avoiding the need for direct harmonization between datasets. For the ROSMAP cohort, we focused on the DLPFC region, integrating matched transcriptomics, DNA methylation, and proteomics data collected from the same brain region. For MSBB, we utilized RNA-seq data from four brain regions (BA10, BA22, BA36, BA44), integrating these with available proteomics and methylation profiles. Because the data preprocessing and multiomics data integration were performed within each cohort independently using anatomically and experimentally consistent data.

To enhance the informativeness of downstream analyses, only the top 75% of variance for each omics dataset including gene expression, DNA methylation and proteomics collected from brain tissues were retained for both ROSMAP and MSBB.

In addition to the integrative analyses of multi-omics data from brain tissues, we conducted a PheWAS using the short whole-genome sequencing dataset from the All of Us Research Project. Leveraging electronic health records (EHR), we investigated the association between phenotypic information, condensed into International Classification of Diseases (ICD) codes, and genetic variations associated with gene expression, known as eQTLs. Our objective was to determine if the eQTLs of DEGs between the two clusters also showed associations with other phenotypes, particularly those related to cognition.

By focusing on eQTLs linked to DEGs identified in brain tissues[12], we explored associations with various phenotypes to elucidate the broader physiological implications of the identified molecular signatures. This comprehensive analysis aims to enhance our understanding of AD heterogeneity and provide valuable insights into the potential regulatory mechanisms underlying AD pathology.

Together, the integration of multiomics data from both ROSMAP and MSBB, coupled with the comprehensive genomic data from the All of Us Research Project, offers a robust framework for dissecting the molecular underpinnings of AD. By elucidating the intricate interplay between genetic, epigenetic, and proteomic factors, our study contributes to the broader effort of unraveling the complexity of AD and advancing personalized approaches to diagnosis, treatment, and disease management.

## Methods

### AD Patient Clustering with Subspace Merging Algorithm

To integrate heterogeneous omics data for patient stratification in Alzheimer’s Disease (AD), we adopt a subspace merging algorithm originally developed for integrative cancer analysis [13] with minor modifications to identify AD subtypes with matched three types of omics data collected from AD patient in the ROSMAP study [14]. This method follows the intermediate data integration paradigm by converting each omics dataset into a patient-to-patient similarity graph, performing spectral embedding to obtain subspaces, and subsequently merging them on a Grassmann manifold. This framework captures both shared and omics-specific structure while preserving the local geometry of each data layer. An overview of this approach is shown in Figure.

Let 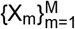 be M omics data matrices, each of dimension N × p_m_ , where N is the number of patients and p_m_ the number of features in the m-th omics layer. Each omics matrix X_m_ was independently standardized to have zero mean and unit variance prior to graph construction and subspace embedding by equation (1), where f is any feature, 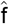 is the corresponding normalized feature, E(f) is the sample mean and the Var(f) is the sample variance for the selected feature. . This normalization ensured comparable scaling within each modality and cohort while preserving cohort-specific biological variation.

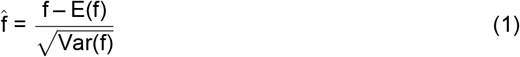

For each omics modality, we construct a similarity graph G^(m)^ by first computing the pairwise heat kernel similarity matrix with equation(2), where*σ*_i_ = distance to 7th nearest neighbor of x_i_ We then define the pairwise similarity matrix S^(m)^ ϵ R^N*×*N^ using a symmetric heat kernel. This kernel formulation guarantees that patients with similar profiles form locally dense neighborhoods, while preserving relative scaling across modalities. We then compute the normalized graph Laplacian L^(m)^ ϵ R^N*×*N^ for each similarity matrix S^(m)^ with equation(3), where D^(m)^ is a diagonal matrix with 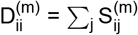. This corresponds to the symmetric normalized Laplacian.

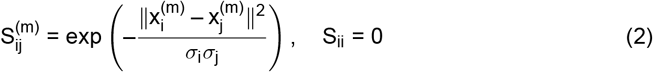

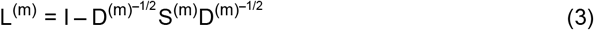

We then perform eigen decomposition of L^(m)^, and obtain the k eigenvectors corresponding to the smallest eigenvalues. These eigenvectors U^(m)^ ϵ R^N*×*k^ form the subspace representations for each modality. Each U^(m)^ is a point on the Grassmann manifold 𝒢(k, N), which denotes the collection of all k-dimensional subspaces in R^N^. The Grassmann manifold provides a principled geometric framework for modeling subspaces, enabling us to quantify the dissimilarity between modality-specific patient embeddings.

We measure the dissimilarity between two subspaces using the squared projection Frobenius norm:

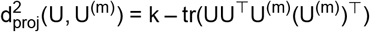

This projection metric quantifies the distance between the projection matrices of U and U^(m)^, capturing how aligned the patient representations from different modalities are in terms of their subspace orientation. To construct a consensus representation U, we jointly minimize two objectives: the spectral loss from each omics graph and the projection distance between U and the modality-specific embeddings. The resulting optimization problem is by solving equation(4):

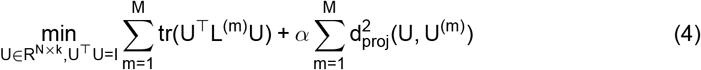

The first term in equation(4) encourages U to preserve the local manifold structure encoded in each Laplacian L^(m)^, while the second aligns the consensus with the individual subspaces. The hyperparameter *α* controls the balance between structure preservation and modality agreement. This leads to the following closed-form solution:

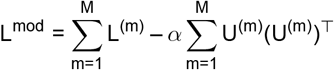

We then obtain the final merged subspace U by computing the top k eigenvectors of L^mod^. Unlike iterative optimization methods, this eigendecomposition-based approach is computationally efficient and converges to a global solution. Our framework treats all omics layers with equal importance during integration. Both the Laplacians and the subspace projection terms are summed without scalar weighting. This reflects an assumption of equal biological contribution across gene expression, proteomics, and DNA methylation.

The rows of the merged subspace U provide patient-level embeddings that capture integrative structure across omics. We apply K-means clustering to U to define patient subtypes. To determine the optimal number of clusters k, we evaluate the average silhouette score for a range of candidate values (e.g., k = 2 to 8). For each value of k, we normalize the eigenvectors of L^mod^, perform K-means clustering with multiple restarts (n = 10), and compute the silhouette score using the Euclidean distance. The value of k that yields the highest average silhouette score is selected as the optimal cluster number. This procedure ensures that the final clustering structure is both geometrically coherent and supported by internal validation metrics.

### Differential Expression Analysis

Differentially gene expression and downstream pathway analysis were applied to establish the gene signatures between the subtypes (patient clusters) for both ROSMAP and MSBB, as well as to the transcriptomic data of the peripheral blood monocyte (PBMC) from the matching patients in ROSMAP in search for possible blood markers. Differential expression analysis was conducted using the linear modeling approach implemented in the ‘limma’ package. The batch effect was corrected and confounding variables such as age and sex were adjusted if the processed data had not applied such adjustment. The empirical Bayes method implemented in ‘limma’ was used to moderate standard errors and obtain more stable estimates of gene-wise variances.

Contrasts were defined to test the difference between the two subgroups identified from the previous steps. The resulting differential expression statistics, including log-fold changes and moderated t-statistics, were used to identify genes that were significantly differentially expressed. To control the false discovery rate (FDR), the Benjamini-Hochberg(BH) procedure was applied to adjust p-values for multiple testing. Genes with an adjusted p-value below a specified threshold (FDR < 0.05) were considered statistically significant.

### Enrichment Analysis on DEGs

DEGs were further analyzed for biological relevance and functional enrichment using ‘clusterProfiler’ package[15], which performs Gene Ontology (GO) and pathway enrichment analysis by applying a hyper-geometric test to identify significantly enriched GO terms and pathways among a set of genes based on their annotations, identifying over-represented biological processes, molecular functions, or cellular components.

### PheWAS

A PheWAS is a type of analysis that explores the association between genetic variants and a wide range of phenotypic traits or outcomes across an entire population or cohort. Unlike traditional genome-wide association studies (GWAS), which typically focus on a specific disease or trait, PheWAS examines the genetic influences on diverse phenotypes, including diseases, clinical measurements, laboratory values, and other health-related characteristics.

This approach allows for the identification of genetic variants that may influence various aspects of health and disease beyond those initially considered in the study. PheWAS can uncover unexpected relationships between genetic variants and phenotypes, provide insights into the pleiotropic effects of genes, and contribute to a more comprehensive understanding of the genetic basis of complex traits and diseases.

#### Genotypic Data

We employed genotypic data of the 105 eQTLs of interest from 11,024 unrelated participants, aged 45 to 95, who were diagnosed with AD, diffuse Lewy body disease, behavioral disturbances concurrent with late-onset Alzheimer’s dementia, or had a family history of dementia (parents, siblings, or self) as recorded in their EHR.

#### Phenotypic Data

Phenotype data for each participant were extracted from their electronic health record (EHR) and mapped to phecodes using the R package “PheWAS” [16]. In the phenotype table, each participant was assigned a case/control label for each health outcome. A participant was classified as ‘TRUE’ for a condition if they had at least two distinct instances of the corresponding phecode, indicating they were a case. Conversely, a ‘FALSE’ classification indicated the absence of the health condition, classifying them as a control. Phecodes with less than 20 cases or controls were excluded. And sex-specific phecodes were assigned only to the corresponding sex. In total, 1858 phenotypes were generated for the downstream association test.

#### Statistical Analysis

We performed logistic regression analyses for each phecode, where the binary case-control outcome was regressed on the genotype of each eQTL. The models were adjusted for age, sex at birth, and the first six principal components to correct for ancestry prediction. Multiple testing correction strategies were employed: We applied the Benjamini-Hochberg (BH) method [17] across all association tests and additionally conducted an eQTL-specific correction for tests involving each respective eQTL. An arbitary p-value threshold of 10^−4^ was utilized to capture any potential important associations.

## Results

### Analysis of the ROSMAP Dataset

The algorithm separated these 79 patients from the ROSMAP study into 2 clusters (33 patients in cluster 1 and 46 patients in cluster 2 (Figure 1A) according to their similarities based on three types of omics data: gene expression, proteomics, and DNA methylation expression.

**Figure 1.**
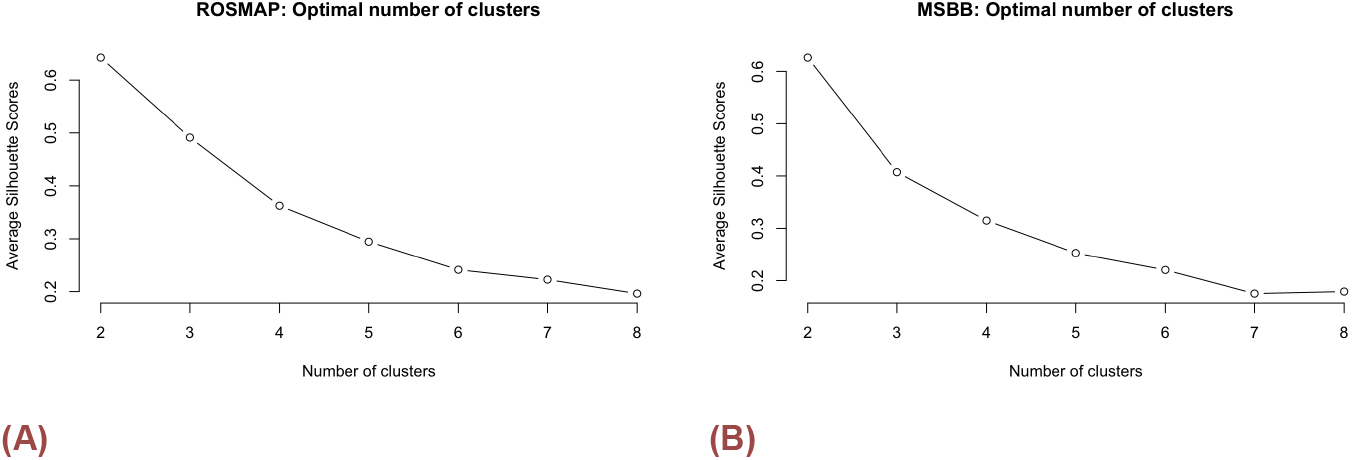
The silhouette plot: for all k’s between 2 and 8, the silhouette score is maximized at k=2 for both A) ROSMAP and B) MSBB, indicating two clusters are well defined, considering both the cohesion within clusters and the separation between clusters

Demographic and diagnostic variables including age at first AD was given (age), sex, Apolipoprotein E (APOE) genotype [18], Clinical diagnosis of cognitive status (dcfdx), Clinical consensus diagnosis of cognitive status at time of death (cogdx), Braak stage score (braaksc) [19], which is a semi-quantitative measure of neurofibrillary triangle pathology, CERAD score (ceradsc), which is a semi-quantitative measure of neuritic plaques were compared between the the two clusters. In comparison, Clinical diagnosis of cognitive status (p value = 0.03598), CERAD score (p value = 0.03548) were significantly different between the two clusters indicating the there were mild differences in disease progression in terms of cognition. And overall, patients in Cluster 1 presented relatively higher cognition score compared with the other Cluster. However, there were no significant difference between age at first Alzheimer’s dementia was given (p value= 0.6463), sex at birth(p value = 1), APOE genotype (p value = 0.7726), Clinical consensus diagnosis of cognitive status at time of death (p value = 0.08646), and Braak stage (p value=0.6157). The subtyping reveals the difference that cannot be explained by age, sex or APOE status, they are revealed only by integrating the multi-omics data-driven approach.

Differential expression analyses were independently conducted for the three omics data types. In proteomics, 87 features were found to be differentially expressed at a significance level of FDR = 0.05. However, the effect size (logFC) between the two clusters was less than 0.2, indicating that the differences were minimal and not biologically substantial (Figure2A). In contrast, none of the other omics data types, including DNA methylation (Figure2B)or RNASeq gene expression data from blood samples (Figure2C), identified any genes with significant differential expression (FDR < 0.05) and a log fold change greater than 0.7. However, a different pattern emerged in the gene expression data obtained from brain samples. In this case, 524 genes exhibited significant differential expression between the two clusters, including 508 upregulated genes in cluster 1 compared to cluster 2, and 16 down-regulated genes(Figure2D).

**Figure 2.**
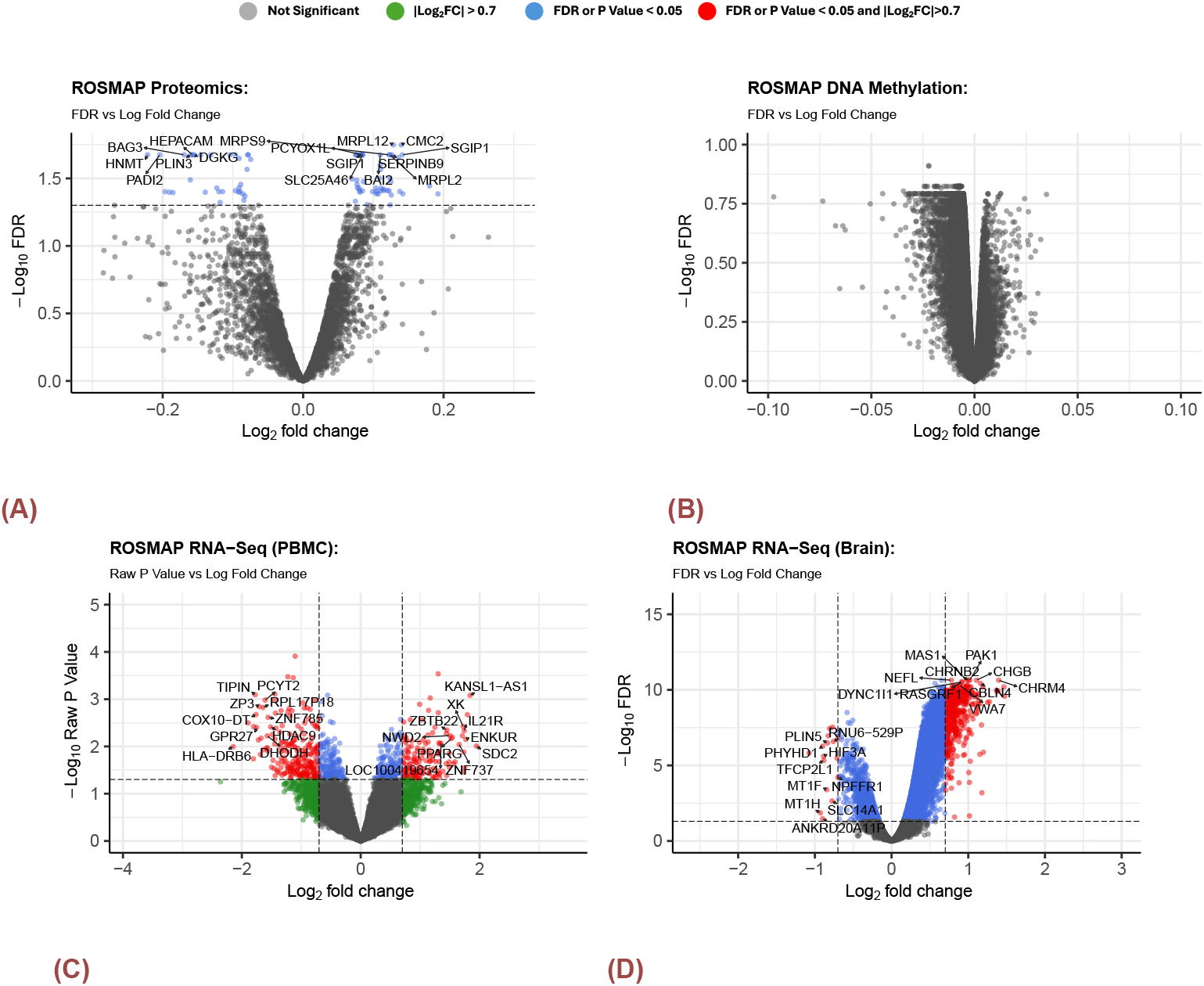
Volcano Plots for DEGs in ROSMAP. A) Bulk Brain RNA-SEq B) Proteomics C) DNA Methylation D)PBMC RNA-Seq. There were no significantly expressed genes or probes for DNA methylation and proteomics data set. There were 524 genes that were significantly expressed between the two clusters. There were 508 up-regulated(cluster1 - cluster2) and 16 down-regulated genes with the magnitude of log fold change greater than 0.7 and FDR less than 0.05. For the RNA-seq collected from PBMC samples, there were 371 genes of interest with the magnitude of log fold change greater 0.7 and raw p-value less than 0.05. Within these genes, 224 were down-regulated and 147 were up-regulated. The top 10 up-regulated and down-regulated genes were labeled with gene symbol in the volcano plot.

Furthermore, there were 154 biological process (BP) terms in Gene Ontology (GO) enriched in the gene set of DEGs identified from the previous section. The examination revealed notable enrichment in categories associated with “synapse organization” (adj.p-value < 0.001) and “cognition” (adj.p-value < 0.001). Figure3A shows a compilation of the top enriched GO BP terms, offering more insight into the key molecular processes associated with DEGs between the two subtypes identified by the subspace merging algorithm. These findings imply that the selected genes are pivotal in brain synaptic activities, cognition, and learning offering a deeper understanding on the molecular mechanisms underlying AD progression.

**Figure 3.**
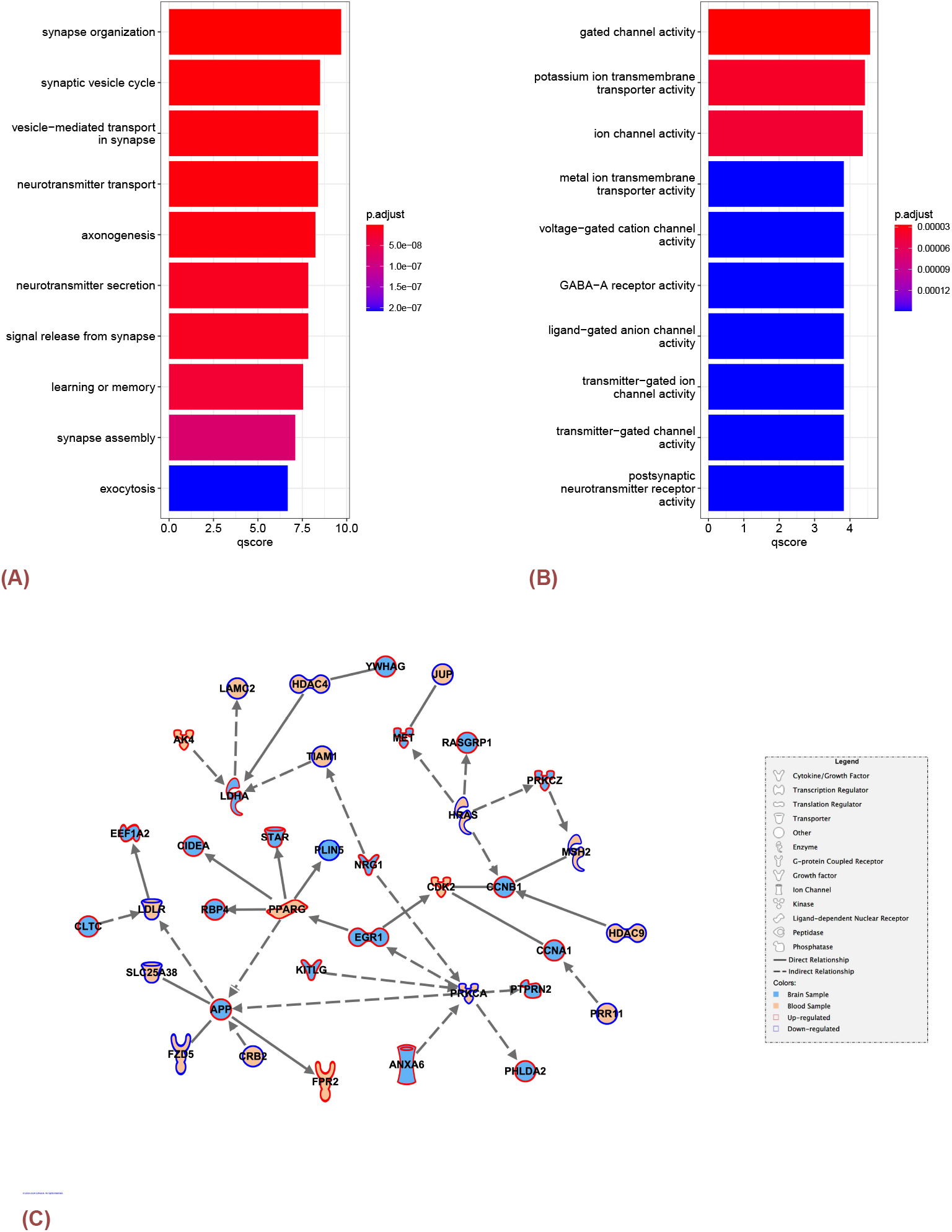
A) The top 10 significantly enriched biological processes on DEGs between the two clusters identified by the subspace merging algorithm in the previous section. B) The top 10 significantly enriched molecular functions for DEGs from brain samples in ROSMAP study. The p-values were corrected with BH method and the qscore was calculated by – log_10_p.adjust.C)The network plot displays the shortest pathways between DEGs in brain tissues and DEGs in blood samples. The shape of the node indicates the type of gene, the edge indicates the direction and type of the relationship between the two connected genes, the fill color indicates whether the gene is expressed in brain or blood, and the outline color indicates whether the gene is up-regulated or down-regulated.

DEGs (both up- and down-regulated genes) from brain samples were used to construct gene sets for enrichment analysis with R package “clusterProfiler”[15]. The analysis revealed significant enrichment in 62 molecular function (MF) terms. For instance, the gene set demonstrated a notable over-representation in terms related to “gated channel activity” (adj.p-value < 0.001) and “potassium ion trans-membrane transporter activity” (adj.p-value < 0.001), and “GABA-A receptor activity” (adj.p-value < 0.001). These findings suggest that the genes differentially expressed between the two subtypes play crucial roles in transport of ions across cell membranes, and forming channels with the ability to open and close in response to specific signals, providing valuable insights into the potential roles of those genes in cellular processes involving ion transport and signaling. Further details and statistical significance measures for each pathway can be referenced in Figure3B.

Despite the absence of genes exhibiting highly statistically significant expression changes with a False Discovery Rate (FDR) less than 0.05, a subset of genes characterized by a relatively substantial effect size (| log FC | ≥ 0.7) has been identified from the PBMC samples. Our analysis involved the systematic exploration and visualization of potential pathways linking blood samples and brain samples. This was achieved by investigating the shortest pathways between all DEGs identified within two clusters from the brain region (comprising 524 genes) and genes exhibiting a large effect size in blood samples (totaling 371 genes). The results revealed networking between multiple genes from blood samples and DEGs within the brain sample (Figure3C).

In the network plot generated through the use of QIAGEN IPA (*QIAGEN Inc*., https://digitalinsights.qiagen.com/IPA)[20], our attention was particularly drawn to the LDLR gene, a member of the Low-Density Lipoprotein (LDL) family. Notably, APOE *ϵ*4 (APOE4) mutation stands as a significant genetic risk factor for AD. Previous studies [21] revealed that APOE plays a crucial role in cholesterol metabolism by binding to various receptors, with LDLR exhibiting a notably high affinity for APOE. The study suggested that it was noteworthy that LDLR was the sole member of its receptor family to demonstrate an isoform-specific binding affinity E4 binds more strongly than E3, which in turn binds more strongly than E2). The LDLR gene regulates ApoE levels in both the periphery and the central nervous system in mouse. It has been identified on astrocytes for this function and was shown to modulate amyloid deposition in AD transgenic mice[22]. Notably, increased LDLR expression level has been associated with reduced amyloid burden and enhanced glial response, suggesting its potential as a therapeutic target[23].

Additionally, we identified two other DEGs with established links to AD: APP and PPARG. The *β*-amyloid precursor protein (APP) gene, which is cleaved to form *β*-amyloid (A*β*), plays a crucial role in the aggregation of A*β* in the brain, a hallmark of AD[24]. Previous literature also reported that APP mRNA is highly expressed in neurons and is upregulated in AD brains, with expression patterns and regulatory transcription mechanisms changing progressively with age and increasingly shifting toward dysfunction.[25]. Furthermore, the nuclear receptor peroxisome proliferator-activated receptor gamma (PPAR*γ*) has emerged as a promising therapeutic target. PPAR*γ* agonists, which regulate glucose and lipid metabolism and suppress inflammatory gene expression, have shown significant improvements in memory and cognition in AD patients, highlighting their therapeutic potential in AD treatment[26].

### Analysis of the MSBB Dataset

The algorithm clustered these 83 patients from the ROSMAP study into 2 clusters (36 patients in cluster 1 and 47 patients in cluster 2 (Figure 1B) according to their similarities based on the following omics data: gene expression for BM10, BM22, BM36, BM44, proteomics, and DNA methylation expression.

We also examined the disparities in demographic and clinical diagnostic variables between the two clusters in the MSBB study. It was found that age at death (p value = 0.607), race (p value = 0.521), sex (p value = 0.660), and APOE genotype (p value = 0.909) did not exhibit significant differences between the two clusters. Conversely, there were significant discrepancies in variables such as A*β* plaque levels (p value = 0.024), Clinical Dementia Rating (CDR) (p value = 0.0474), and Braak stage (p value = 0.021) between the clusters identified through the integration of three types of omics data.

The analysis of gene expression patterns revealed prominent distinctions between clusters identified by the subspace merging algorithm across different brain regions. In the BM10 brain region, 21 genes (5 down-regulated and 16 up-regulated in Figure4A) exhibited statistically significant differential expression level between the identified clusters with FDR < 0.05 and magnitude of logFC ≥ 0.7. Similarly, the BM22 region displayed significant expression differences in 134 genes (24 down-regulated and 110 up-regulated in Figure4B). In BM36, 169 genes (96 down-regulated and 73 up-regulated in Figure4C) showed significant expression changes, and in the BM44 region, 110 genes (32 down-regulated and 78 up-regulated in Figure4D) demonstrated differential expression patterns, indicating discerning regional variations. However, differentally expressed features identified in DNA methylation data (Figure4E). In addition, three up-reuglated genes were significantly expressed in the proteomics data (Figure4F), where A4_Abeta42 referred to as Amyloid-beta protein 42 or Abeta42, is a 42-amino acid peptide derived from the amyloid-beta precursor protein (APP), which differentially expressed in the brain tissue from ROSMAP study as well.

**Figure 4.**
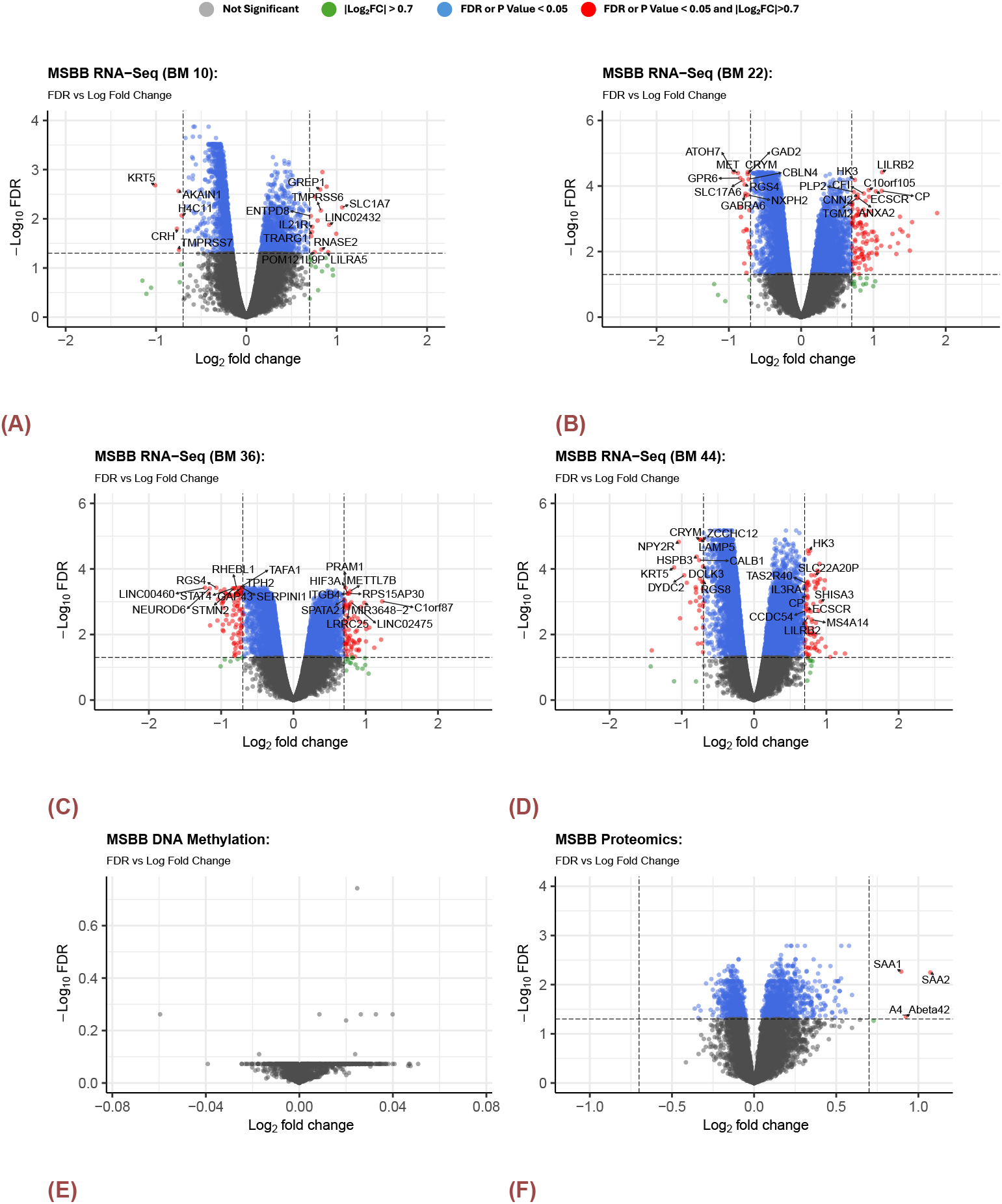
Volcano Plots for DEGs in MSBB by different BM regions. A) RNA-SEq (BM10) B) RNA-SEq (BM22) C) RNA-SEq (BM36) D)RNA-SEq (BM44) E) DNA Methylation F) Proteomics. There were no significantly expressed genes or probes for DNA methylation. 3 DEGs identified from proteomics data.

Furthermore, an exploration of gene intersections between these regions unveiled shared and region-specific expression profiles (Figure5). Notably, only gene SLC1A7, which is a member of the solute carrier (SLC) super-family that specifically encodes the EAAT5 protein, which influence the levels of intracellular chloride within neurons and membrane potential[27], displayed differential expression consistently across BM10, BM22, BM36, and BM44. The number of DEGs in each region and can be found in Figure 5D.

**Figure 5.**
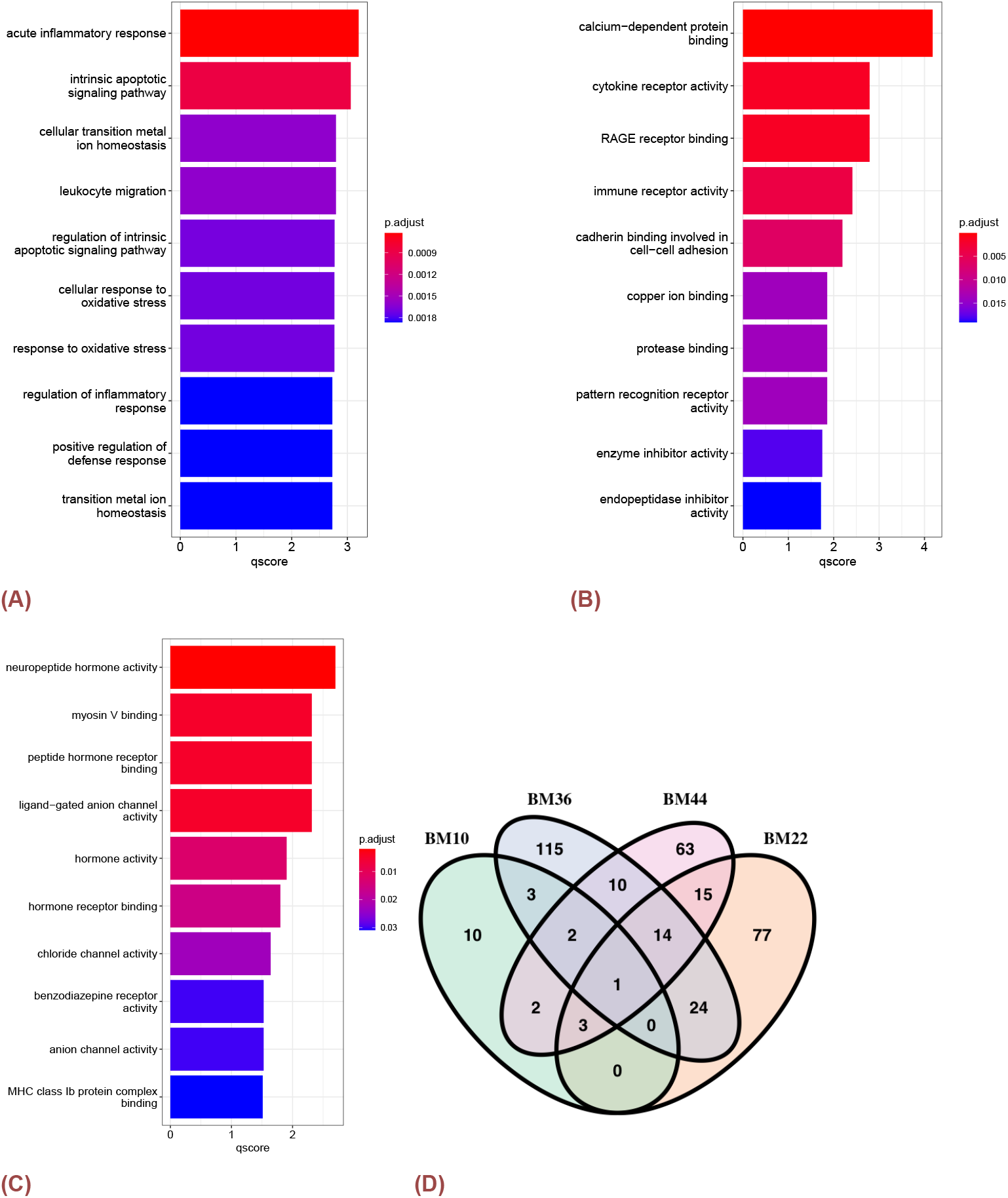
Pathway Analysis for MSBB RNA-Seq data from different brain regions. A)Top 10 significant Biological Processes of DEGs in BM22 B)Top 10 significant Molecular Functions of DEGs in BM22 C)Top 10 significant Molecular Functions of DEGs in BM36 D) the number of DEGs in each region. The p-values were corrected with BH method and the qscore was calculated by – log_10_ p.adjust.

We then conducted enrichment analysis on DEGs for each distinct brain region including BM10, BM22, BM36, and BM44, where multi-testing was adjusted with BH method. The outcomes reveal significant enrichment in diverse biological processes with 108 statistically significant GO terms and pathways, including the acute inflammatory response (adj.p-value < 0.001), intrinsic apoptotic signaling pathway (adj.p-value < 0.001), cellular transition metal ion homeostasis (adj.p-value < 0.001), and leukocyte migration (adj.p-value < 0.001), within the BM22 region (Figure5A). Moreover, 36 significant molecular functions such as calcium-dependent protein binding (adj.p-value < 0.001), cytokine receptor activity (adj.p-value = 0.0016), RAGE receptor binding (adj.p-value = 0.0016), and immune receptor activity (adj.p-value = 0.0038) were identified as enriched in BM22 (Figure5B). These findings collectively suggest a comprehensive molecular landscape, shedding light on the potential functional implications associated with the DEGs within the BM22 brain region. In addition, the BM36 (Figure5C) region exhibited enrichment in 19 statistically significant molecular functions. Noteworthy functions included neuropeptide hormone activity (adj.p-value = 0.0020), myosin V binding (adj.p-value = 0.0048), and peptide hormone receptor binding (adj.p-value = 0.0048). Furthermore, three molecular function terms demonstrated statistical significance in both the BM22 and BM36 regions, namely ligand-gated anion channel activity (adj.p-value = 0.043 in BM22 and adj.p-value = 0.005 in BM36), L-glutamate transmembrane transporter activity (adj.p-value = 0.029 in BM22 and adj.p-value = 0.035 in BM36), and acidic amino acid transmembrane transporter activity (adj.p-value = 0.036 in BM22 and adj.p-value = 0.037 in BM22). These results enhance our comprehension of the intricate molecular profile specific to the BM22 and BM36 regions, and can emphasize commonalities in molecular functional aspects shared between BM22 and BM36 (Figure5).

### PheWAS

While our analysis did not reveal any significant associations after a stringent multiple-testing correction using all association tests, we however identified 16 important associations at the p-value threshold of 10^−4^, which disclose the potential association between pre-selected eQTLs and the Phecodes available in the EHR. The PheWAS results can be found in Table 2. In addtion, within these 16 important associations, 7 associations were significant under the eQTL-specific multi-testing correction strategy.

**Table 2.** PheWAS Results Table: However, we identified 16 important associations at an arbitrary p-value threshold of 10^−4^ suggesting a potential association between 105 pre-selected eQTLs and the Phecodes available in the EHR. The adj.P.Values were adjusted with BH procedure, and the eQTL.P.Values were adjusted with BH method using only associations involving the selected eQTL.

| eQTL | Chr | Gene | Phecode | Phenotype Description | Category | OR | P Value | adj.P Value | eQTL.P.Value |
| --- | --- | --- | --- | --- | --- | --- | --- | --- | --- |
| rs1786605 | 18 | DTNA | 442.4 | Arterial dissection | Circulatory system | 3.570 | $2.273 \times 10^{-5}$ | 0.567 | 0.031 |
| rs1540063 | 18 | DTNA | 442.4 | Arterial dissection | Circulatory system | 3.552 | $2.483 \times 10^{-5}$ | 0.567 | 0.033 |
| rs1540064 | 18 | DTNA | 442.4 | Arterial dissection | Circulatory system | 3.543 | $2.597 \times 10^{-5}$ | 0.567 | 0.035 |
| rs4808761 | 19 | RAB3A | 530.3 | Stricture and stenosis of esophagus | Digestive | 1.764 | $2.656 \times 10^{-5}$ | 0.567 | 0.036 |
| rs11678354 | 2 | MEIS1 | 292.1 | Aphasia/speech disturbance | Mental Disorders | 0.595 | $2.980 \times 10^{-5}$ | 0.567 | 0.040 |
| rs11670315 | 19 | RAB3A | 530.3 | Stricture and stenosis of esophagus | Digestive | 1.756 | $3.023 \times 10^{-5}$ | 0.567 | 0.041 |
| rs11681729 | 2 | MEIS1 | 292.1 | Aphasia/speech disturbance | Mental Disorders | 0.596 | $3.299 \times 10^{-5}$ | 0.567 | 0.044 |
| rs9489197 | 6 | ROS1 | 963.1 | Antineoplastic and immunosuppressive drugs causing adverse effects | Injuries and Poisonings | 3.167 | $4.153 \times 10^{-5}$ | 0.567 | 0.056 |
| rs9320603 | 6 | ROS1 | 963.1 | Antineoplastic and immunosuppressive drugs causing adverse effects | Injuries and Poisonings | 3.167 | $4.162 \times 10^{-5}$ | 0.567 | 0.056 |
| rs9320602 | 6 | ROS1 | 963.1 | Antineoplastic and immunosuppressive drugs causing adverse effects | Injuries and Poisonings | 3.166 | $4.185 \times 10^{-5}$ | 0.567 | 0.056 |
| rs11086095 | 19 | RAB3A | 530.3 | Stricture and stenosis of esophagus | Digestive | 1.736 | $4.424 \times 10^{-5}$ | 0.567 | 0.060 |
| rs6707386 | 2 | PAX8-AS1 | 870.3 | Other open wound of head and face | Digestive | 2.856 | $6.910 \times 10^{-5}$ | 0.686 | 0.090 |
| rs9789607 | 2 | MEIS1 | 292.1 | Aphasia/speech disturbance | Mental Disorders | 0.612 | $7.120 \times 10^{-5}$ | 0.686 | 0.096 |
| rs75271568 | 7 | PTPRN2 | 288 | Diseases of white blood cells | Hematopoietic | 0.503 | $7.575 \times 10^{-5}$ | 0.686 | 0.102 |
| rs4849177 | 2 | PAX8-AS1 | 870.3 | Other open wound of head and face | Injuries and Poisonings | 2.806 | $9.480 \times 10^{-5}$ | 0.686 | 0.125 |
| rs6755040 | 2 | PAX8-AS1 | 870.3 | Other open wound of head and face | Injuries and Poisonings | 2.805 | $9.520 \times 10^{-5}$ | 0.686 | 0.124 |

Notably,the two eQTLs of RAB3A, which was a DEG from the brain tissues: rs11670315 (OR = 1.756,p-value = 3.023×10^−5^, Figure 6A) and rs4808761 (OR = 1.764, p-value = 2.656×10^−5^, Figure 6B) on chromosome 19 were strongly associated with phecode 530.3 (Stricture and stenosis of esophagus) in the digestive category. In addition, the two eQTLs of MEIS1 that was identified from the blood sample: rs11681729 (OR = 0.596,p-value = 3.299×10^−5^, Figure 6C) and rs11678354 (OR = 0.595, p-value = 2.980×10^−5^, Figure 6D) on chromosome 2 were strongly associated with phecode 292.1 (Aphasia/speech disturbance), which was in the mental disorders category.

**Figure 6.**
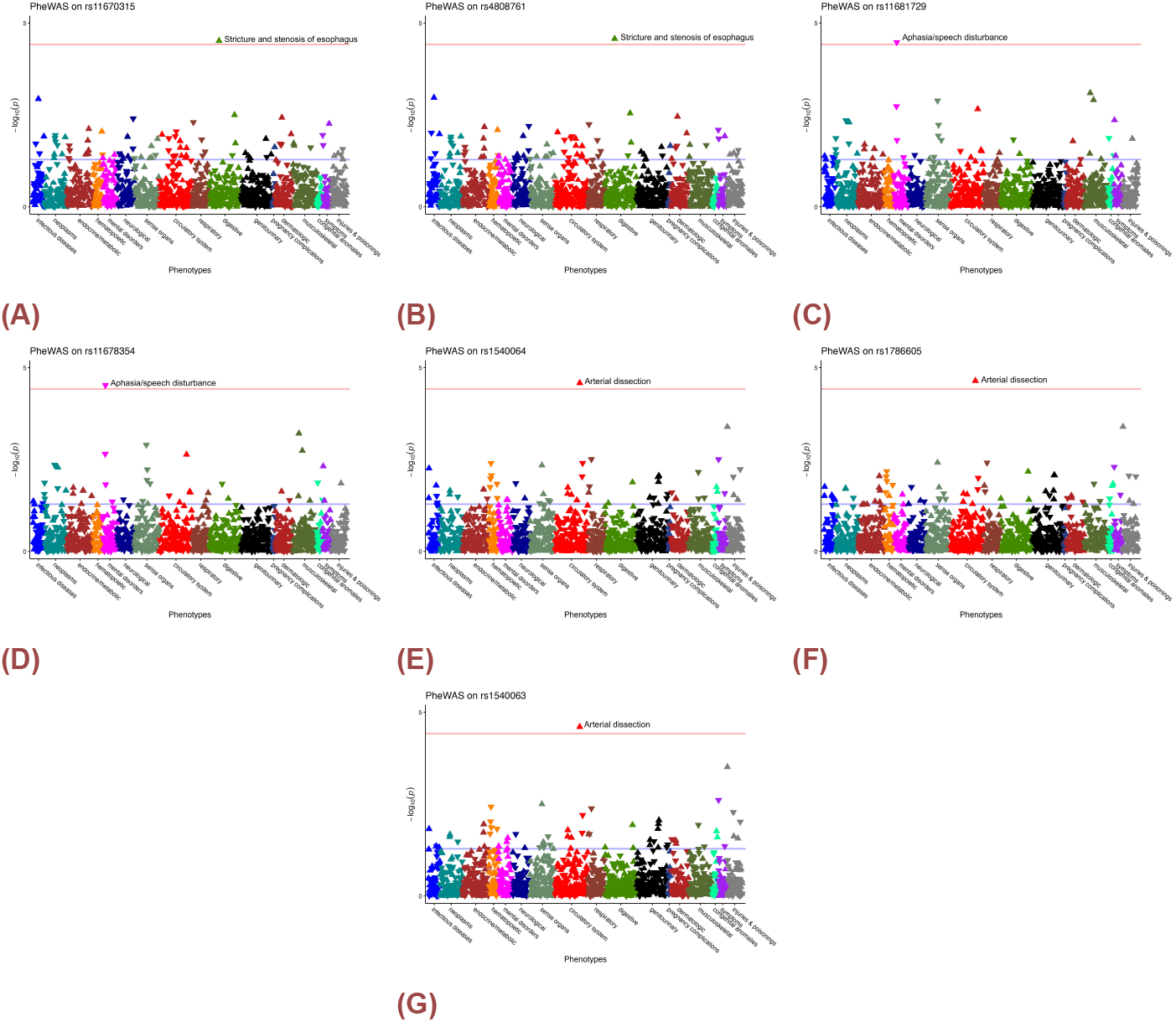
Manhattan plot of eQTLs. The y-axis presents the – log_10_ raw p-value and the x-axis present the phenotype category. The arrow presents the direction of the odd ratio. The blue line suggested raw p-value at level 0.05, and the red line present the Bonferroni corrected p-value using only all association tests involving the selected eQTL. A) rs11670315, B) rs4808761, C) rs11681729, D) rs11678354, E) rs1540064, F) rs1786605 and G) rs1540063.

Furthermore, the three eQTLs of DTNA, which was a DEG from blood sample also: rs1540064 (OR = 3.542,p-value = 2.597×10^−5^, Figure 6E), rs1786605 (OR = 3.570,p-value = 2.273×10^−5^, Figure 6F), and rs1540063 (OR = 3.552, p-value = 2.482×10^−5^, Figure 6G) on chromosome 18 were strongly associated with phecode 442.4 (Arterial dissection) in the circulatory system.

## Discussion and Conclusions

The Subspace Merging Algorithm was able to integrate multiple types of omics data and separate AD patients from multiple cohorts into clusters with distinctive cognitive and brain pathology. The resulted clusters do not show significant differences in age, gender, or APOE status and are revealed only by data-driven integrative multi-omic analysis for both ROSMAP and MSBB studies.

The low-density lipoprotein receptor gene (LDLR) showed a larger effect size in comparing the expression level between the two subtypes identified by the subspace merging algorithm. Preceding pre-clinical experiments have strongly suggested that LDLR enhances the clearance of brain amyloid-beta (A*β*), thereby modulating A*β* metabolism. LDLR emerges as a pivotal pathway influencing A*β* metabolism, rendering it an attractive therapeutic target for AD, as indicated by Kim et al. (2009) [28]. Consequently, the interaction between APOE and LDLR holds the potential to significantly impact the risk and progression of AD. Paradoxically, our analysis revealed that LDLR expression was lower in Cluster 1, which exhibited better cognitive outcomes, contrary to intuitive expectations based on its purported protective role against AD pathology. Consequently, the interaction between APOE and LDLR holds the potential to significantly impact the risk and progression of AD. A comprehensive understanding of the underlying biological mechanisms of this functional interaction can contribute to more effective treatment design and management strategies for different subtypes of AD.

The results supported that the gene LDLR could be a novel blood biomarker for identifying AD subtypes. The subspace merging algorithm integrated multiple types of omics data from the same cohort of patients and successfully identified two clusters within AD patients. Differential expression analysis identified 524 significant genes from the brain samples and 371 genes displaying large effect size from the PBMC samples. The network analysis indicated that the LDLR gene could be a novel blood biomarker for identifying AD subtypes. Future follow-up work may focus on validating whether the LDLR gene could act as a subtype identification marker in other publicly accessible datasets with similar study design, and whether these biological alternations could be explained clinically.

In addition, we identified novel association between genetic variation of DEGs between the two clusters and common phenotypes information available in EHR in utilizing genotypic and phenotyping information collected from All of Us research workbench. Despite not finding significant associations after multiple-testing corrections, our analysis identified 16 notable associations at a p-value threshold of 10^−4^. And 7 signficant associations were revealed with eQTL-specific multi-testing correction, including the potential association between eQTLs of MEIS1, a risk factor of sudden cardiac death and Aphasia/speech disturbance, the association between eQTL of DTNA with Arterial dissection, a circulatory system disorder, and the association between eQTLs of RAB3A, which was down-regulated in AD patients, with a digestive disorder named stricture and stenosis of esophagus.

These findings highlighted potential genetic links between these eQTLs and other diseases, warranting further investigation into their biological mechanisms and potential clinical implications. Our PheWAS results also provided insights into novel association between genetic variation and clinical phenotyping information, such as the alternative allele of both rs11678354(T/A) and rs11681729 (A/G) was associated with an increase in gene expression level of MEIS1, a gene had been identified as a risk factor for sudden cardiac death and previous mice model suggested that MEIS1 interacts with various trophic factors signaling pathways during postmitotic neurons differentiation[29]. Further both these two eQTLs were negatively associated with the risk of Aphasia/speech disturbance, indicating as the number of alternative allele appeared in patient’s genotypic profile, the risk of getting the disease would decrease.

Furthermore, the alternative allele of both rs11670315(G/A) and rs4808761 (A/C) was associated with an increase in gene expression level of RAB3A, which was up-regulated in the cluster with higher cognitive test scores in ROSMAP dataset. Previous studies identified this gene could regulate A*β* production, and was deregulated in AD brains [30, 31, 32, 33]. In addition, the PheWAS results suggested that with an increase in the number of alternative alleles of both eQTLs, the risk of stricture and stenosis of esophagus would increase.

Last but not least, the expression level of DTNA was associated with the frequency of alternative allele of the three eQTLs: rs1540064 (A / G), rs1540063 (C / G), and rs1786605 (G / A). Previous studies revealed that the expression profile of DTNA in the hippocampus (HIP) increased significantly among subjects with dementia and associated levels of phosphorylated tau (P-tau) in the temporal cortex[34]. And a meta-analysis of gene expression data also revealed that this gene was up-regulated in three studies on CA1 of the HIP of brains with AD[35]. The PheWAS results indicated that as the frequency of alternative alleles in these eQTLs increases, the risk of arterial dissection also increases, enriching our understanding of how specific genetic variations could influence multiple health conditions, especially those related to cognition and AD pathology.

To summarize, in this study we applied advanced graph-based unsupervised machine learning algorithm to characterize potential subtypes in AD using multi-omics data from brain samples. We were able to identify different groups of AD patients with different clinical presentations and further identified potential blood-based markers and explored the association of the phenotypes with genetic variants of these marker genes.

While our subspace merging framework provides a structured approach to integrating multiomics data for identifying AD subtypes, several limitations merit discussion. First, our analysis relies exclusively on postmortem brain tissue, which may not fully capture the dynamic and temporal nature of AD progression. Second, although two large-scale cohorts (ROSMAP and MSBB) were included, external validation was not conducted due to anatomical and technical heterogeneity between datasets, limiting the generalizability of the identified subtypes.

Third, the integrative model assumes equal contribution from each omics layer. While this assumption simplifies computation and reduces overfitting risk, it may obscure modality-specific signals that are biologically relevant to certain pathways or patient subgroups. Additionally, as is common with unsupervised clustering, the identified subtypes require further validation in prospective or clinical settings, particularly in the context of treatment response and disease progression.

Future directions include incorporating longitudinal or in vivo data such as imaging or blood-based biomarkers. Adaptive fusion strategies—such as modality-specific weighting or attention mechanisms—could enhance the model’s ability to prioritize more informative data layers. While we initiated joint modeling of brain and peripheral tissues, extending this approach through transfer learning or multitask learning may support more clinically actionable subtyping. With the rapid progress in machine learning, we will also explore advanced graph-learning methods to further refine subtype discovery and representation.

## Resource availability

### Lead contact

Ziyan Song

### Materials availability

This study did not generate new unique reagents.

### Data and code availability

The code supporting the findings of this study is openly available at Github, located at https://github.com/ziyansong08/Identify-AD-subtypes-and-markers-multiomic-data-subspace-merging-algorithm.git.

Data for the ROSMAP are available through the AMP-AD Knowledge Portal: RNA-Seq (Brain) syn8691134, RNA-Seq (PBMC) syn22024496, Proteomics syn17015098, DNA Methylation syn3157275, clinical information syn3191087, and eQTL analysis results syn16984409.

Data for the MSBB are available through the AMP-AD Knowledge Portal: RNA-Seq syn7391833, Proteomics syn24995077, DNA Methylation syn21447661, and clinical information syn6101474.

This study used data from the All of Us Research Program’s Controlled Tier Dataset V7, available to authorized users on the Researcher Workbench.

### Data Preprocessing

#### The ROSMAP Dataset

##### DNA Methylation Data

The DNA methylation profiles of each patient were generated utilizing the Illumina Human-Methylation450 beadset. To address missing values, a k_th_-nearest neighbor algorithm (k=100) was applied for imputation. Following this, adjustments were made for age, sex, and experimental batch[10]. Subsequent analyses included only probes located on the CpG island and those associated with promoters or cell type-specific promoters.

##### Transcriptomic Data

Bulk RNA-seq data sequenced from DLPFC samples were collected from AD knowledge portal. The normalized log2(CPM) of RNA-seq expression values of samples collected from DLPFC regions were adjusted for confounding variables including PMI, sex, and age at death[36, 37]. The limma package[38] was used to correct the sequencing batch.

Monocyte RNA-seq data was sequenced on NovaSeq 6000. Pre-processing involved trimming for adapter sequences using Cutadapt tools, followed by quality control checks encompassing adapter content, per-sequence, and per-base quality, GC content, and sequence duplication levels. Alignment to the hg38 human reference genome was performed using the STAR aligner, with a subsequent generation of a count table of uniquely mapped reads from genes utilizing the HTseq-count tool from the HTseq package[39].

##### Proteomic Data

The proteomics data was labeled using a mass spectrometry-based protein quantification approach with tandem mass tag (TMT)[40, 41, 42]. Furthermore, the “limma” package was used to mitigate the batch effects and the effect of confounding variables such as age at death and sex. The reads were then median-centered and log_2_transformed.

##### eQTL Data

To elucidate the complex relationships between genetic variation, gene expression, and phenotyping outcomes, we collected eQTL data from ROSMAP[12] via AD Knowledge Portal. This dataset included SNP-gene associations with corresponding p-values and effect sizes. For each DEG, we selected only the top three significant eQTLs with an FDR less than 10^−4^ to prevent over-representation. This selection process yielded a total of 105 eQTLs derived from 37 DEGs, 49 from blood DEGs and 56 from brain DEGs, for subsequent analysis.

#### The MSBB Dataset

##### DNA Methylation

For the DNA methylation array data, genomic DNA was extracted from 10mg of postmortem brain tissue from the parahippocampal gyrus and hybridized onto Infinium MethylEPIC BeadChips. Raw array data from IDAT files were preprocessed and normalized by R package “minfi”[43], methylation levels at each CpG site on the Illumina 850K platform were represented by *β* values, and further adjustments for batch effect and confounding variables including age at death, sex and race were made accordingly using limma package.

##### Transcriptomic Data

The RNA-seq data were collected from Brodmann Areas 10, 22, 36, and 44 and processed by aligning raw sequence reads to the human genome hg19 with the STAR aligner (v2.3.0e)[44]. Gene-level expression was then quantified using featureCounts (v1.4.4) from the Subread package [45], retaining only genes with a read count of at least 1 in a minimum of 10 libraries. Normalization was performed using the trimmed mean of M-values (TMM) method in the “edgeR” [46] R package to account for differences in library size, and correction for known covariate factors including PMI, race, batch, sex, RIN and exonic rate was applied to mitigate confounding effects.

##### Proteomic Data

Protein abundances in postmortem cortical tissues obtained from Brodmann area 36 of the parahippocampal gyrus were quantified using the TMT technique, with subsequent normalization, correction for batch effects, and adjustments for confounding variables such as sex, age at death, and race.

#### All of Us Dataset

##### Genotypic Data

In the All of Us Research Program, short whole-genome sequencing data, the genotypic information, were obtained by processing DNA samples collected from participants who consented to participate in the program and donated fresh whole blood (4 ml EDTA and 10 ml EDTA) as the primary source of DNA. The short whole-genome sequencing data were processed following standardized laboratory protocols such that bar-coded libraries were constructed for PCR-free sequencing on the Illumina NovaSeq 6000 instrument and sophisticated quality control procedures were implemented at various stages of the sequencing process. Initial quality control involved utilizing the Illumina DRAGEN pipeline, which assessed metrics such as contamination levels and mapping quality. Furthermore, joint calling across the dataset enhanced sensitivity and identified samples warranting further scrutiny due to potential issues[47].

## Acknowledgement

This project is partially supported by the NIH 5U54AG065181 (Indiana TREAT-AD Center) and NIH R21AG075541 (Indiana University Indiana University Precision Health Initiative).

We extend our sincere gratitude to the participants of the All of Us Research Program. We also acknowledge the National Institutes of Health’s All of Us Research Program for providing access to the participant data examined in this research.

The data available in the AD Knowledge Portal would not be possible without the participation of research volunteers and the contribution of data by collaborating researchers. The results published here are in whole or in part based on data obtained from the AD Knowledge Portal (https://adknowledgeportal.org).

Study data were provided by the Rush Alzheimer’s Disease Center, Rush University Medical Center, Chicago. Data collection was supported through funding by NIA grants P30AG10161 (ROS), R01AG15819 (ROSMAP; genomics and RNAseq), R01AG17917 (MAP), R01AG30146, R01AG36042 (5hC methylation, ATACseq), RC2AG036547 (H3K9Ac), R01AG36836 (RNAseq), R01AG48015 (monocyte RNAseq) RF1AG57473 (single nucleus RNAseq), U01AG32984 (genomic and whole exome sequencing), U01AG46152 (ROSMAP AMP-AD, targeted proteomics), U01AG46161(TMT proteomics), U01AG61356 (whole genome sequencing, targeted proteomics, ROSMAP AMP-AD), the Illinois Department of Public Health (ROSMAP), and the Translational Genomics Research Institute (genomic). Additional phenotypic data can be requested at www.radc.rush.edu.

The MSBB data were generated from postmortem brain tissue collected through the Mount Sinai VA Medical Center Brain Bank and were provided by Dr. Eric Schadt from Mount Sinai School of Medicine.

The MSBB proteomics data were provided by Dr. Levey from Emory University based on postmortem brain tissue collected through the Mount Sinai VA Medical Center Brain Bank provided by Dr. Eric Schadt from Mount Sinai School of Medicine.

Study data were provided through the Accelerating Medicine Partnership for AD (U01AG046161 and U01AG061357) based on samples provided by the Rush Alzheimer’s Disease Center, Rush University Medical Center, Chicago. Data collection was supported through funding by NIA grants P30AG10161, R01AG15819, R01AG17917, R01AG30146, R01AG36836, U01AG32984, U01AG46152, the Illinois Department of Public Health, and the Translational Genomics Research Institute.

Data generation was supported by the following NIH grants: P30AG10161, P30AG72975, R01AG15819, R01AG17917, R01AG036836, U01AG46152, U01AG61356, U01AG046139, P50 AG016574, R01 AG032990, U01AG046139, R01AG018023, U01AG006576, U01AG006786, R01AG025711, R01AG017216, R01AG003949, R01NS080820, U24NS072026, P30AG19610, U01AG046170, RF1AG057440, and U24AG061340, and the Cure PSP, Mayo and Michael J Fox foundations, Arizona Department of Health Services and the Arizona Biomedical Research Commission.

We thank the participants of the Religious Order Study and Memory and Aging projects for the generous donation, the Sun Health Research Institute Brain and Body Donation Program, the Mayo Clinic Brain Bank, and the Mount Sinai/JJ Peters VA Medical Center NIH Brain and Tissue Repository. Data and analysis contributing investigators include Nilüfer Ertekin-Taner, Steven Younkin (Mayo Clinic, Jacksonville, FL), Todd Golde (University of Florida), Nathan Price (Institute for Systems Biology), David Bennett, Christopher Gaiteri (Rush University), Philip De Jager (Columbia University), Bin Zhang, Eric Schadt, Michelle Ehrlich, Vahram Haroutunian, Sam Gandy (Icahn School of Medicine at Mount Sinai), Koichi Iijima (National Center for Geriatrics and Gerontology, Japan), Scott Noggle (New York Stem Cell Foundation), Lara Mangravite (Sage Bionetworks).

## Author contributions

Conceptualization, Z.S., K.H. and J.Z.; Methodology, Z.S., X.H., K.H., and J.Z.; Visualization, Z.S.; Formal Analysis, Z.S., and A.J.J; Data Curation, Z.S. and X.H.; Writing – Original Draft, Z.S. and K.H.; Writing – Review & Editing, Z.S., X.H., A.J.J., T.S.J, K.H., and J.Z.; Funding Acquisition, K.H. and J.Z.; Project Administration, K.H. and J.Z.; Supervision, K.H. and J.Z..

